# Inferring the Joint Distribution of Structural and Functional Connectivity in the Human Brain using UNIT-DDPM

**DOI:** 10.1101/2024.02.03.578750

**Authors:** Varun Canamedi

## Abstract

The structural wiring of the brain is expected to produce a repertoire of functional networks, across time, context, individuals and vice versa. Therefore, a method to infer the joint distribution of structural and functional connectomes would be of immense value. However, existing approaches only provide deterministic snapshots of the structure-function relationship. Here we use an unpaired image translation method, UNIT-DDPM, that infers a joint distribution of structural and functional connectomes. Our approach allows estimates of variability of function for a given structure and vice versa. Furthermore, we found a significant improvement in prediction accuracy among individual brain networks, implicating a tighter coupling of structure and function than previously understood. Also, our approach has the ad-vantage of not relying on paired samples for training. This novel approach provides a means for identifying regions of consistent structure-function coupling.

## 1 Introduction

For most biological systems function is highly determined by the underlying anatomical structure. The structural substrate of the brain consists of a collection of neuronal populations interconnected by a complex anatomical network referred to as the connectome (1). The connectome strongly constrains neural activity by inducing correlated activation and deactivation patterns commonly referred to as functional networks (2), thereby influencing processes such as perception, cognition, and behavior (3). Understanding the interplay between brain structure and function is an important goal in neuroscience (4).

Recent advances in multi-modal brain imaging have enabled us to study this relationship in detail. Tractography algorithms applied on diffusion-weighted MRI can be used to infer structural connectivity (5), (6), while similar techniques can be used to derive functional connectivity from functional MRI (7). All these factors have resulted in a growing emphasis on conceptualizing structure-function relationship in the nervous system as associations between structural and functional brain networks.

Numerous studies have confirmed that functional connectivity is the strongest in structurally connected regions. Although this is true, recent evidence suggests that the structure-function relationship is more nuanced than a simple linear correspondence (8), as the brain can form functional connections even in the absence of structural links (9). Higher order interactions between neural elements may be influenced by polysynaptic communication between multiple structural elements, giving rise to this complex relationship (10).

By accounting for these higher order interactions, various approaches such as biophysical models (11), (12), (13), network communication models (14), (15), graph harmonics (16) and multivariate statistical models (17), (18) have provided evidence of a more accurate modelling of the structure-function relationship compared to univariate models. These methods can be used to simulate functional dynamics within a structurally connected population. The functional connectivity predicted by these models can be compared against functional connectomes generated from empirical data. While these mechanistic approaches have considerable advantages, they provide poor generalizations of the SC-FC relationship.

Supervised learning approaches have been proven instrumental in overcoming the limitations of mechanistic models (19), (20), (21), (22). These approaches utilize a large data set of empirical structural and functional connectome pairs to learn the unidirectional mapping between them. Significantly higher prediction accuracies achieved by these approaches have provided compelling evidence to suggest that a tighter coupling between structure and function does exist in nature (19). Furthermore, numerous studies have demonstrated that these approaches can be used to accurately estimate individual connectomes while preserving inter-individual variability. A given structural connectome can be expected to produce a variety of co-activation patterns, across time, context, and individuals (23). However, current approaches can only estimate a single FC ma-trix for a given SC. They do not provide any means for modelling the inherent variability of brain networks. The nature of SC-FC relationship is more nuanced than a unidirectional mapping (24). It is rather a joint distribution of structural and functional connectomes. The SC-FC relationship when represented as a joint distribution is far closer to reality than when represented as a unidirectional mapping.

Thus, compared to learning the unidirectional mapping between the two domains, inferring the joint distribution of structural and functional connectomes would be of immense value in advancing our understanding of the brain as a biological system. The closest existing approaches can get to inferring the joint distribution is by training two models, one predicting functional connectomes with structural connectomes as input and the other predicting structural connectomes from functional connectomes as input. Even then, the deterministic nature of these models would make it impossible to estimate the distribution of the probable outputs for a given input.

In the present study, we a use UNIT-DDPM (25) which is an unsupervised image translation algorithm to model the task of decoding the structure-function relationship in the human brain as an unpaired image translation problem. While all earlier studies consider learning a deterministic unidirectional mapping between structural and functional networks, we demonstrate the effectiveness of the UNIT-DDPM framework for learning the bilateral transform paths between these networks. The stochastic nature of the sampling algorithm produces a collection of longitudinal data series from an inferred joint distribution of structural and functional connectomes for all inputs.

To the best of our knowledge, the present study is one the first to attempt the task inferring the joint distribution of structural and functional connectomes. Also, while reiterating the strong structure-function coupling in the human connectome reported by previous studies, this study provides new evidence to suggest that the percentage of variation among individual networks can be captured more accurately even in the absence of paired examples during training.

## 2 Materials and Methods

### 2.1 Datasets

In this study, we have used the Human Connectome Project (HCP) s1200 dataset (26). The HCP s1200 dataset is a collection of brain imaging data from 1200 healthy young adult participants.

### 2.2 Structural Connectivity Mapping

We utilized the minimally pre-processed diffusion weighted MRI data for 450 individuals from the HCP dataset. The preprocessing steps included corrections for eddy currents, head motion and non-linear distortion. The acquisition process involved capturing 90 gradient directions at three b-values (1000, 2000, and 3000 s/mm2) using spin-echo planar imaging. The MR acquisition protocols and pre-processing steps are described in detail elsewhere. We employed whole-brain fiber tracking on the preprocessed dMRI data. Specifically, we utilized constrained spherical deconvolution (CSD) (27), a method that estimates multiple fiber orientations within each white matter voxel. This information was then utilized to propagate streamlines throughout the white matter volume using a deterministic fiber tracking algorithm.

For the deterministic tractography, we employed CSD-based techniques and uniformly seeded streamlines from a white matter mask derived from automated structural segmentation. To ensure comprehensive coverage, the boundaries of the white matter mask were slightly expanded to address any potential gaps between gray and white matter boundaries. The streamlines were propagated using the sdstream option in the tckgen function of the MRtrix software package, employing default parameters for step size, angle threshold, and FOD (Fiber Orientation Distribution) threshold.

**Figure 1.**
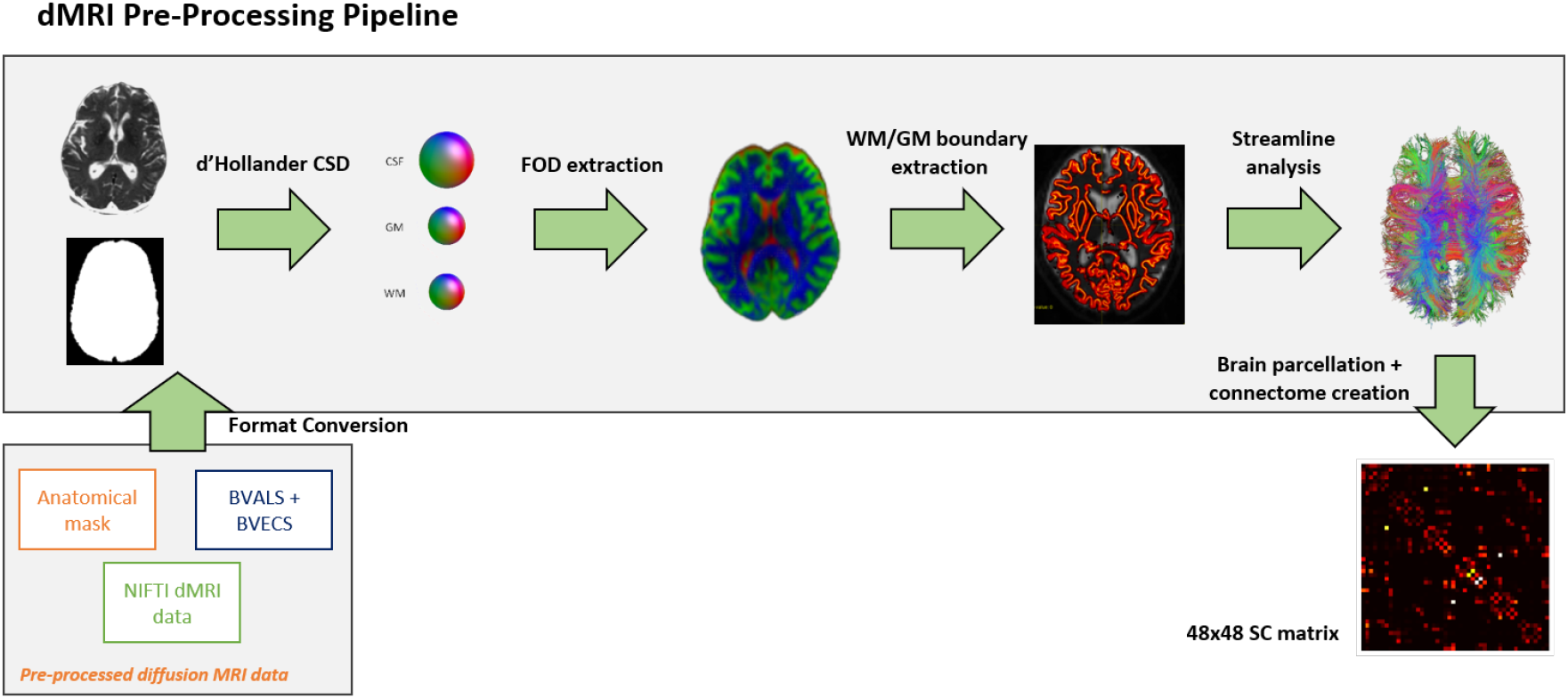
Data pre-processing operations performed on diffusion weighted MRI data for the calculation of structural connectivity matrix (eSC) using Harvard-Oxford sub-cortical parcellation

We set the CSD with a spherical harmonic order of 8, generated up to 1 million streamlines per individual, and limited the maximum streamline length to 400 mm. From these streamlines, we constructed an individual’s structural connectivity matrix, which quantified the number of streamlines connecting each pair of regions in the Harvard-Oxford cortical parcellation atlas. Each element of the matrix represented the structural connectivity strength between two regions, denoting the count of streamlines with one endpoint in region i and another endpoint in region j.

### 2.3 Functional Connectivity Mapping

We obtained minimally pre-processed resting state functional MRI (rs-fMRI) data for 450 individuals from the HCP dataset. The for the UNIT-DDPM model selection of individuals for the training set was performed in a randomized fashion in order to ensure the absence of paired examples. An alternate dataset containing functional MRI data from the same individuals selected for structural connectivity mapping was obtained for create a paired dataset for training supervised models.

The rs-fMRI data acquisition involved employing a multiband gradient-echo planar imaging (EPI) sequence, with a repetition time (TR) of 720 milliseconds, a multiband factor of 8, and subjects instructed to keep their eyes open and maintain relaxed fixation on a bright cross-hair projected against a dark background. Two runs of EPI data were collected, utilizing both right-to-left and left-to-right phase encodings. Minimal pre-processing steps included data cleaning using independent component analysis with a focus on fixing artifacts (ICA-FIX) and nonlinear transformation to the Montreal Neurological Institute (MNI) standard space.

After temporally demeaning the two runs, we concatenated them to create a total of 2400 volumes for each individual. To quantify the strength of functional connectivity, we calculated a functional connectivity matrix for each individual using the Pearson correlation coefficient. Specifically, voxel-specific time series were spatially averaged within each region to generate regionally averaged time series. The Pearson correlation coefficient between the regionally averaged time series of regions of i and j were was stored in element (i, j) of the individual’s functional connectivity matrix. Since these matrices were inferred from actual MRI data, we refer to them as empirical structural connectivity matrices (eSC) and empirical functional connectivity matrices (eFC).

### 2.4 UNIT-DDPM

A UNIT-DDPM model was trained on an unpaired dataset containing 400 structural and functional connectivity matrices sampled from the HCP s1200 dataset. The UNIT-DDPM algorithm used in this study was adopted from Sasaki and colleagues (25). Basically, Denoising Diffusion Probabilistic Models (DDPMs) iteratively destroys structure within a data distribution with increasing gaussian noise. In the reverse process, the algorithm learns to restore data yielding a highly flexible generative model of the data.

**Figure 2.**
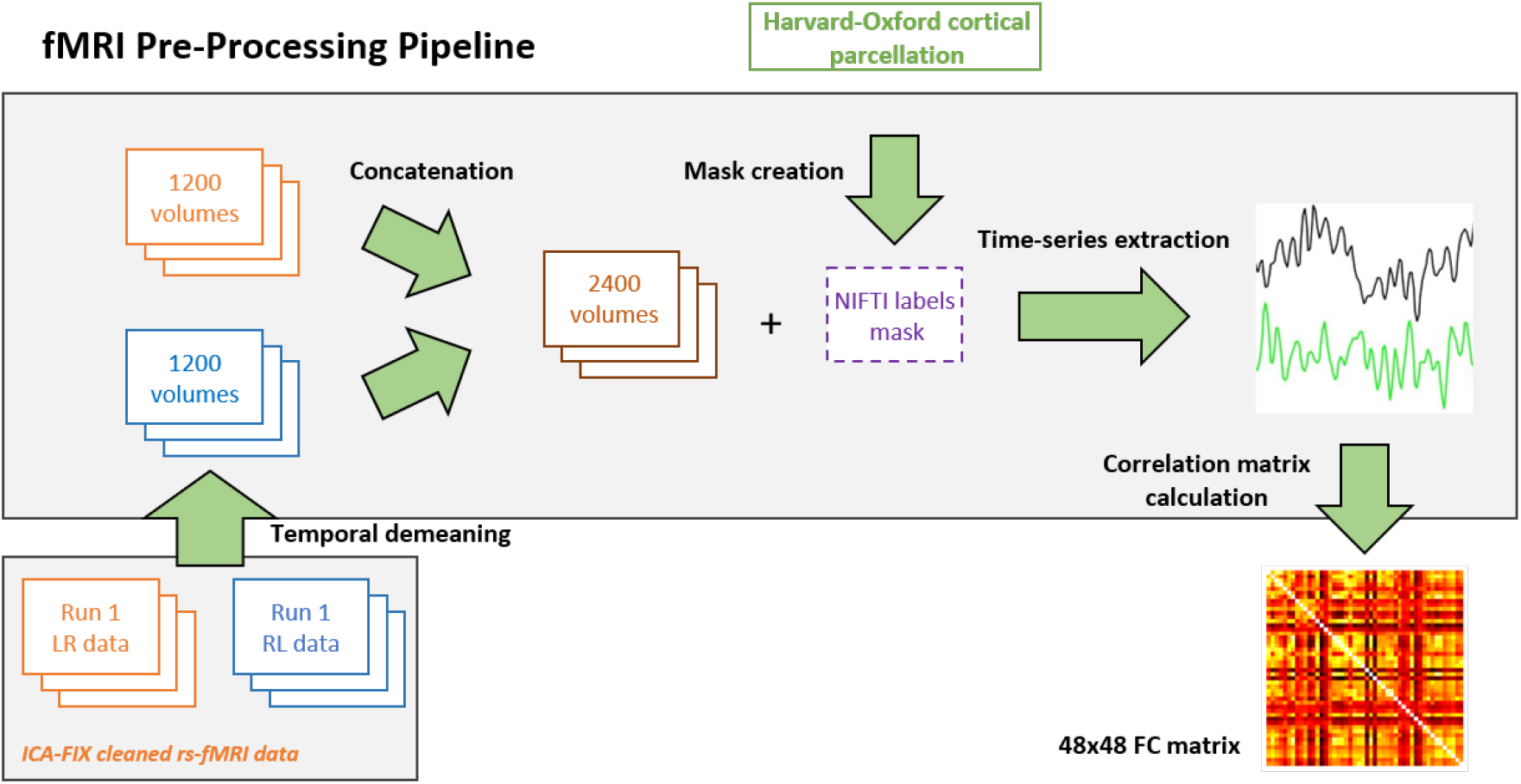
Data pre-processing operations performed on resting state functional MRI data (Left-to-Right and Right-to-Left scans) for calculating functional connectivity matrices using Harvard-Oxford sub-cortical parcellation.

DDPM employs a unique approach to model data by treating it as a latent variable described by p(x0) := p(x0:T)dx1:T. Here, x0 q(x0) represents the initial images, T signifies the length of the Markov chain, and x1, …, xT denote the latent variables, sharing the same dimensional characteristics as the images. Within this framework, p(x0:T) corresponds to a Markov chain equipped with learned Gaussian transitions, constituting the reverse process where:

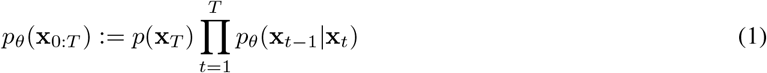

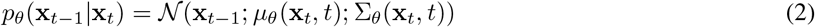

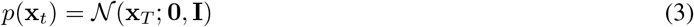

Furthermore, in the forward process, DDPM also approximates the posterior *q*(*x*_1:*T*_| *x*0), wherein a Markov Chain incrementally introduces progressive Gaussian noise to the images:

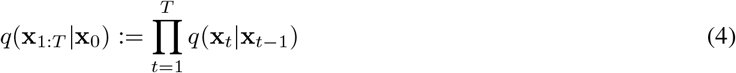

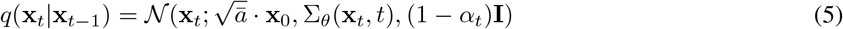

Here, *α*_*t*_ ∈ *α*_1_, …, *α*_*T*_ represents scheduled weights of the gaussian noise. Equation (4) gradually adds noise based on a variance schedule *α*_*t*_. The sampling *x*_*t*_ at any timestep t purely as a function of *x*_0_ as given in the below equation:

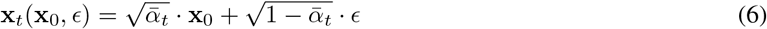

where, 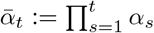 and *ϵ ∼ 𝒩* (0, **I**) by optimizing model parameter *θ* through minimizing the loss function given by,

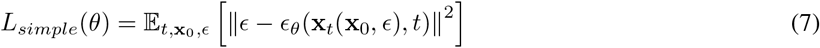

The UNIT-DDPM algorithm uses the latent information estimated by the DDPM to learn various image domains and establish connections between their respective latents. Consequently, it enables the gradual generation of samples starting from noise, progressively refining them into images within the target domain. This process maintains a meaningful relationship with the input source domain images.

Let, 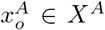 represent the source domain and 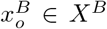 represent the target domain. UNIT-DDPM iteratively optimizes are only utilized during model training for bi-directional translation. In the reverse diffusion optimization step, model parameters are tuned to minimize the loss function which can be reframed as:

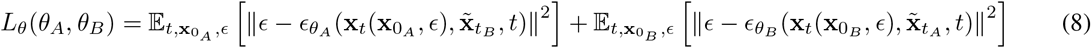

The translation function parameters are also updated to minimize the DSM objective fixing the diffusion model parameters. Where:

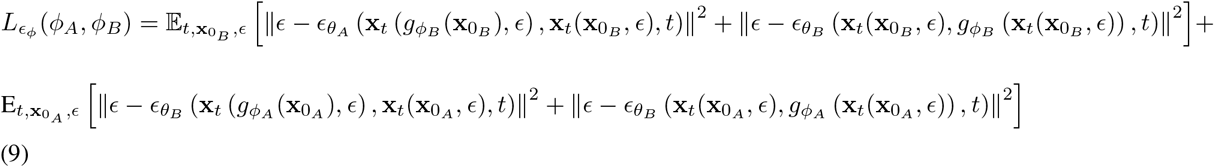

Also, training is regularized by the cycle-consistency loss to make domain translation bi-directional. Thus, the loss function is described as:

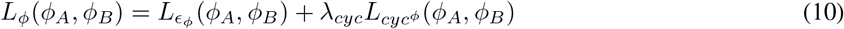

where *λ*_*c*_*yc* is the weight parameter for the cyclic loss function 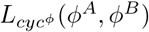

**Figure 3.**
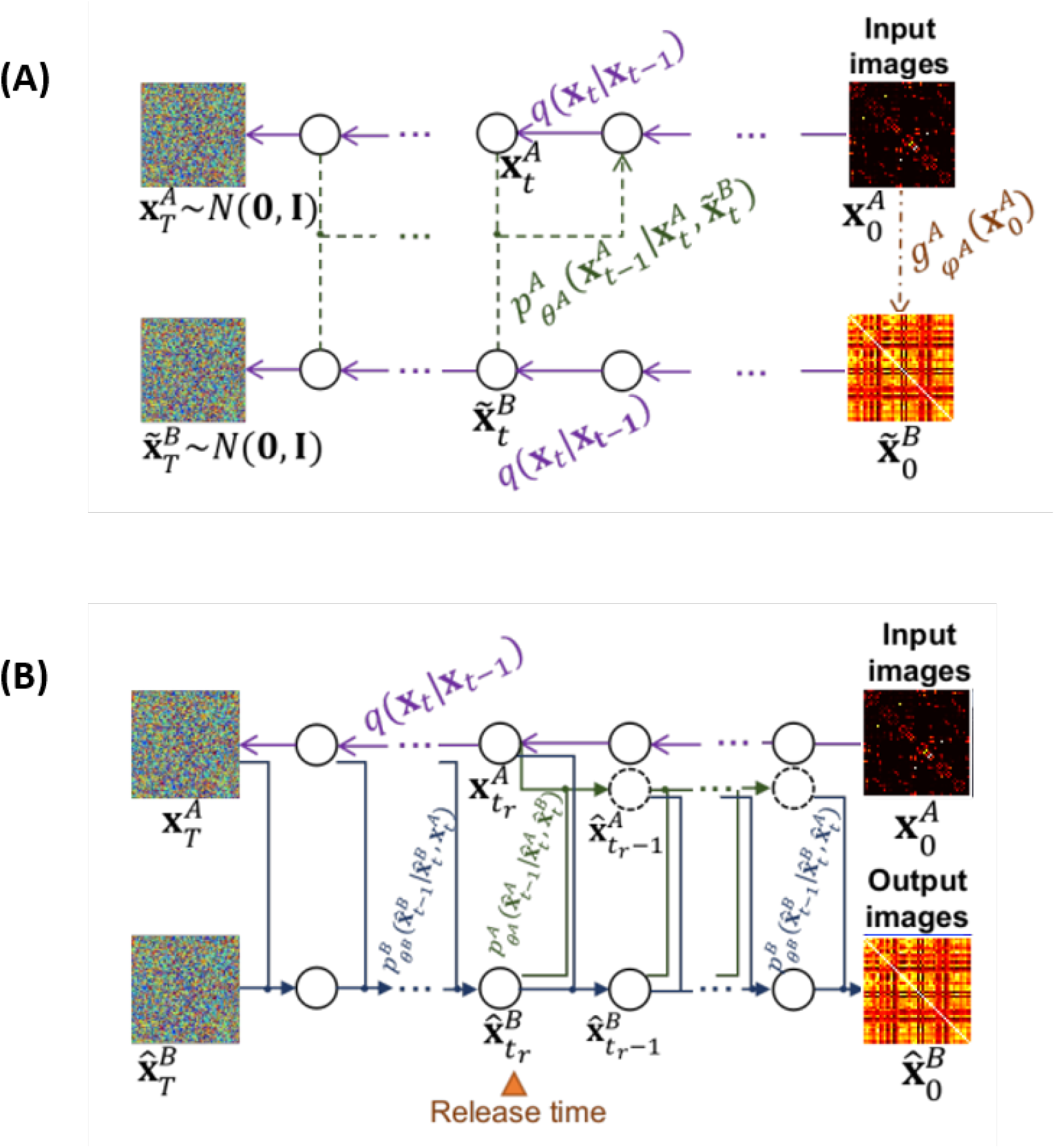
UNIT-DDPM (A) Model Training on unpaired SC and FC matrices and (B) Domain translation (inference) with trained models.

The model parameters *ϕ*^*A*^ and *ϕ*^*B*^ obtained during training are used for translating the input images from source domain to target domain. The domain translation functions are discarded and the target domain images are inferred progressively from white noise. During the sampling phase, the generative process is conditioned on the input source domain images that have undergone perturbation by the forward process starting from t = T and continuing until a selected arbitrary time step *t*_*r*_ within the range [1, T]. Subsequently, these conditioned images are regenerated by the reverse process starting from this specific time step, which we refer to as the ‘release time’

Our method implements the denoising models using an attention U-Net architecture, which are a variant of the U-Net architecture that incorporates attention mechanism as shown in Figure 4. This allows the network to focus on the most important features of the latent representation during the denoising process. For the domain translation functions, we use simple U-Net models. This is a departure from the original UNIT-DDPM, which used simple U-Net models for denoising and ResNet for domain translation functions during training.

**Figure 4.**
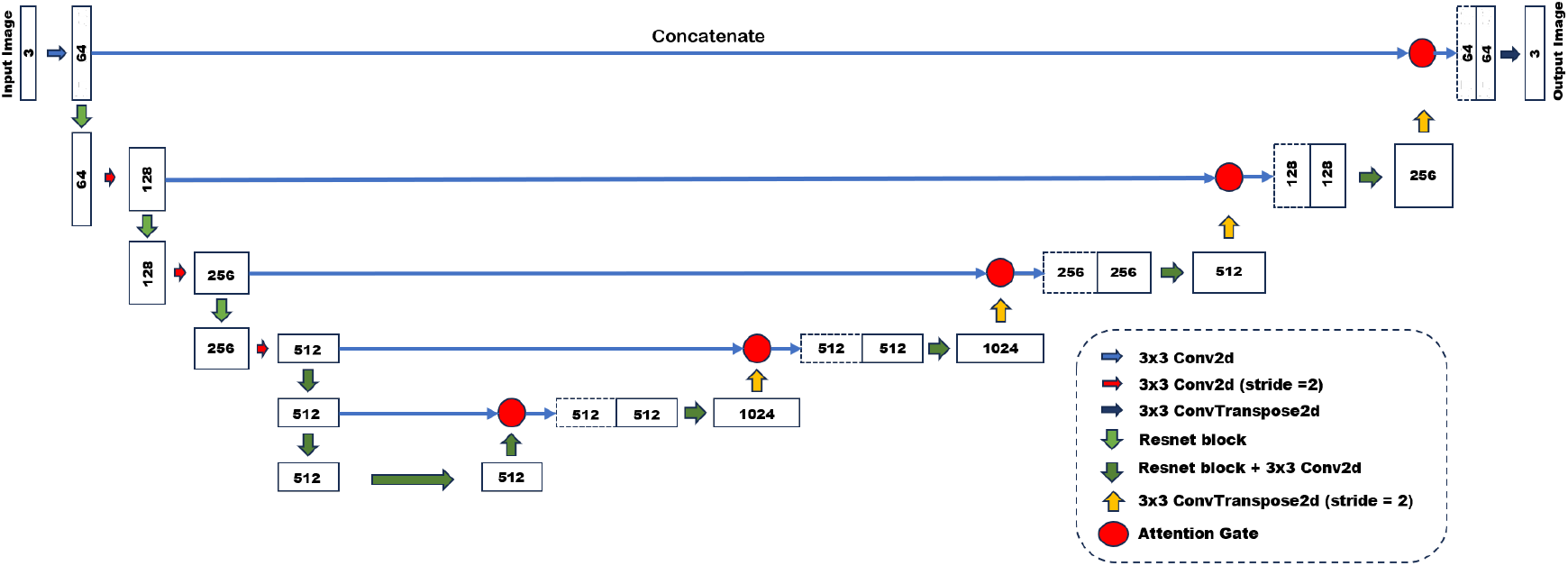
The diagram of the implemented Attention U-net architecture. Each attention gate contains ReLu, sigmoid and resampler block.

### 2.5 Multi-Layer Perceptron

In order to perform a comparative analysis between our method and existing state-of-the-art, we also train a Multi-Layer Perceptron (MLP) model implemented by Sarwar and colleagues (19). The network consisted of 8 fully connected hidden layers with 1024 neurons each with a dropout of 0.5. Two activation functions, namely, leaky ReLU and hyperbolic tangent (tanh) were used alternatively and the Glorot algorithm was used the initialize model weights. The feed-forward network was trained with the Adam optimizer. The network is provided with the rescaled upper-triangular elements of the structural connectivity matrix as input and trained to predict the functional connectivity matrix for each individual. The main objective to the training process is to learn the uni-directional mapping between SC and FC matrices while maximizing the similarity between the predicted matrix and the empirical matrix. In order to address the trade-off between prediction accuracy and inter-individual variability, the loss function of the multi-layer perceptron is given by (11):

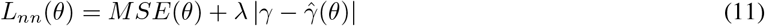

where, *γ*, 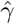 represent empirical inter-individual heterogeneity and heterogeneity among model predictions respectively and *γ* is the regularization constant. The expression for *γ* and 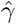 are given by (12) and (13):

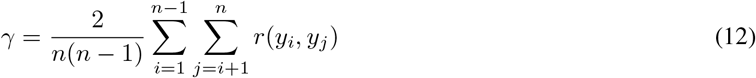

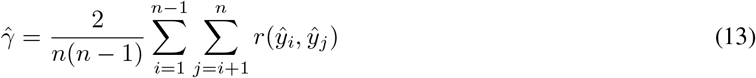

Details regarding hyper-parameter settings and implementation of both the UNIT-DDPM and the multi-layer perceptron is provided in Table 1.

**Table 1:** Hyper-Parameter Settings.

| Parameters | Multi-Layer Perceptron | UNIT-DDPM |
| --- | --- | --- |
| Epochs | 10,000 | 10,000 |
| Batch size | 16 | 16 |
| $\lambda$ | 0.45 | - |
| Training Subjects | 400 (paired) | 400 (unpaired) |
| Test Subjects | 50 | 50 |
| Time steps | - | 999 |

## 3 Results

We mapped whole-brain structural and functional networks for 450 individuals participating in the Human Connectome Project (HCP). Structural and functional connectivity matrices were estimated for 48 regions using the Harvard-Oxford sub-cortical parcellation atlas on both diffusion weighted MRI and resting state functional MRI data. Among the 450 individuals, the SC and FC matrices for 400 individuals were select randomly to ensure the absence of paired examples for model training and 50 paired matrices were used for testing. The UNIT-DDPM was trained on the unpaired dataset to infer the joint distribution of SC and FC.

The functional connectivity (FC) matrices calculated from resting state fMRI are referred to as empirical FC (eFC) and the structural connectivity matrices (SC) calculated from diffusion weighted MRI are referred to as empirical SC (eSC). The functional and structural connectivity matrices estimated by the model on the test set are referred to as pFC and pSC respectively. To assess model performance and quantify the strength of SC-FC coupling, correlation coefficients (pearson-r) were computed between pFC-eFC and pSC-eSC pairs across all regions. In order to evaluate the ability of the model to retain inter-individual variations in its predictions, we computed the correlation between each distinct pair of individuals, referred to as inter-pFC, inter-eFC, inter-pSC and inter-eSC. We also calculated correlation coefficients for eSC-eFC, eSC-pFC, eFC-pSC pairs for each individual in the test set.

In order to benchmark the performance of the UNIT-DDPM model, a state-of-the-art feed-forward Multi-layer Perceptron (MLP) was trained in a supervised manner on an alternate dataset containing paired SC-FC examples to predict functional connectivity from structural connectivity. The hyperparameters for the supervised model were selected as suggested by Sarwar and colleagues. We use pFCNN and pFCdiff nomenclature to differentiate between the predictions of the UNIT-DDPM from the MLP mode.

### 3.1 Prediction of group-averaged structural and functional connectivity

The strength of structure-function coupling is typically quantified using group-averaged connectivity measures. Therefore, structure-function coupling at a group level for both the MLP model as well as UNIT-DDPM. Group-averaged matrices were obtained from the 50 individuals from the test set. The computed eSC and eFC matrices were used by the UNIT-DDPM to predict the pFC_diff_ and pSC_diff_ matrices respectively. The trained MLP model was used to predict FC from SC.

We find that the prominent off-diagonal elements in the FC matrix were better estimated by the MLP by a small margin. Comparison between Figure 6 (A) and Figure 6 (B) shows that pFC_diff_ is sparser compared to pFC_NN_. Figure 6 (C) suggests that the UNIT-DDPM is better at estimating FC compared to SC. One reason for this might be the sparse nature of SC matrices make them more difficult to estimate compared to FC matrices.

### 3.2 Prediction of individual structural and functional connectivity

Having found that group level structure-function coupling observed in UNIT-DDPM is similar to that of methods suggested by prior studies, we considered comparing the performance of both models on a much more challenging task of predicting structural and functional connectivity among individual participants. To address the compromise between inter-individual connectivity and intra-individual connectivity, model hyper-parameter for the MLP were chosen as suggested by (19). This ensures that the model simply does not predict the group-averaged matrix for each individual. Overall, the UNITDDPM was better at preserving inter-individual variations while yielding more accurate predictions of individual functional connectivity matrices compared to the MLP model. The low correlation between pFC matrices of both models with respect to eSC as seen in Figure 5 (A) suggests that they are not simply recapitulating the inputs provided during testing. The almost perfect matching between the means of inter-individual across all unique pairs of individuals between eSC and pSCdiff suggests that UNIT-DDPM is better at retaining heterogeneity among SC matrices compared to FC. The trade-off between inter-individual correlation and intra-individual correlation is clearly evident as an increase in heterogeneity in model outputs leads to a decrease in intra-individual prediction accuracy as seen in Figure 5 (B) and Figure 5 (E).

**Figure 5.**
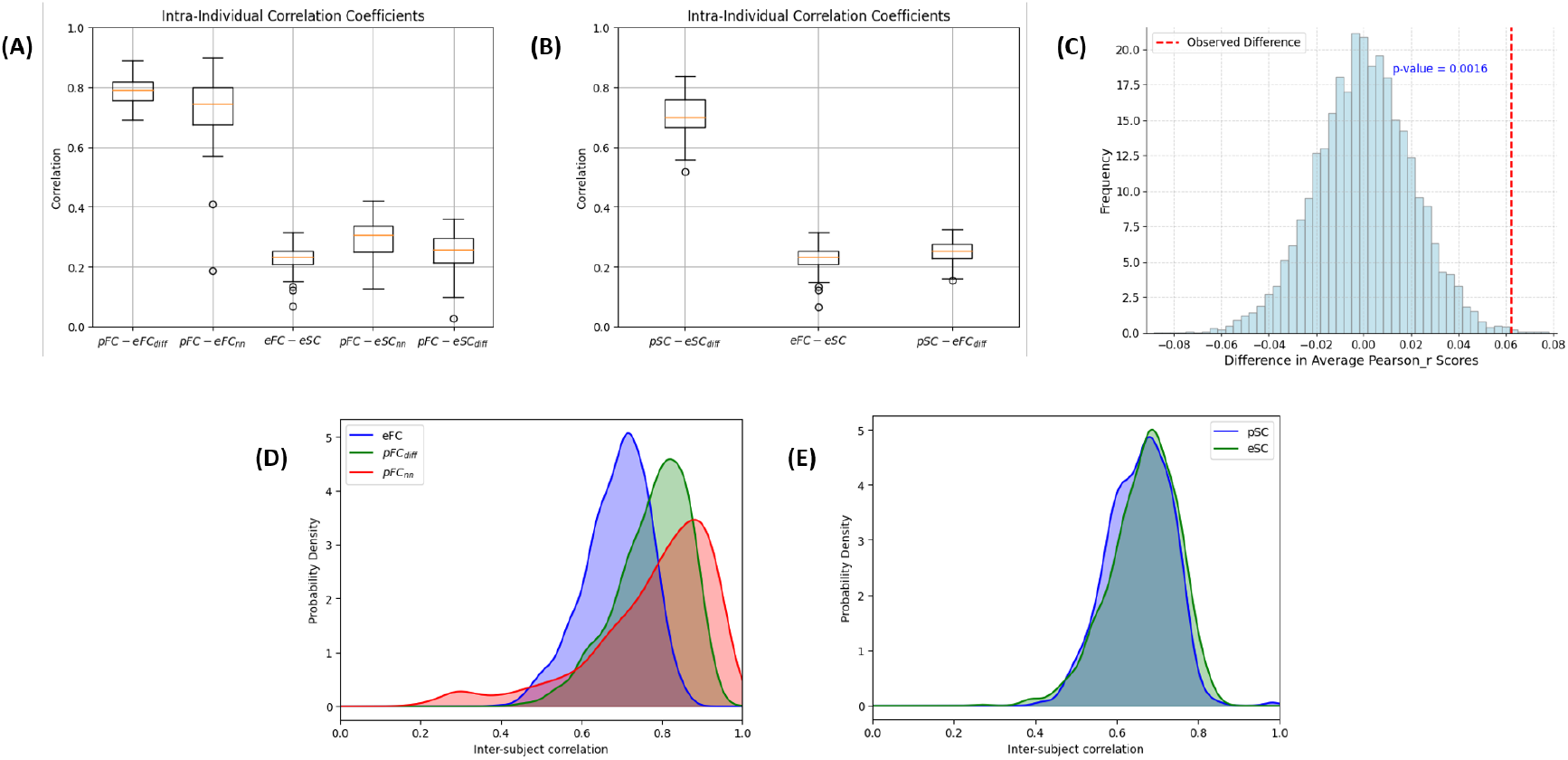
Predicting function from structure and vice-versa in the human connectome based on individual connectivity matrices (N = 50) using UNIT-DDPM and Multi-layer Perceptron. (A) Boxplots show the distribution of correlation coefficients for 50 individuals. The two leftmost boxplots represent the intra-individual correlation between eFC and pFC which is the primary measure of prediction accuracy for both models. The three rightmost boxplots represent correlation with respect to eSC. The orange mark within the boxplots represents the median. (B) The leftmost boxplot represents the distribution of correlation coefficients between eSC and pSC which measures accuracy of the UNIT-DDPM in predicting structural connectivity from functional connectivity. The two rightmost boxplots represent correlation with respect to eFC. (C) The graph represents the bootstrap distribution of difference in average correlation scores between pFC*−* eFC_diff_ and pFC *−*eFC_NN_. (D) and (E) represent the distribution of inter-individual correlation coefficients for all distinct pairs among 50 individuals. Lower inter-individual correlation represents greater heterogeneity.

**Figure 6.**
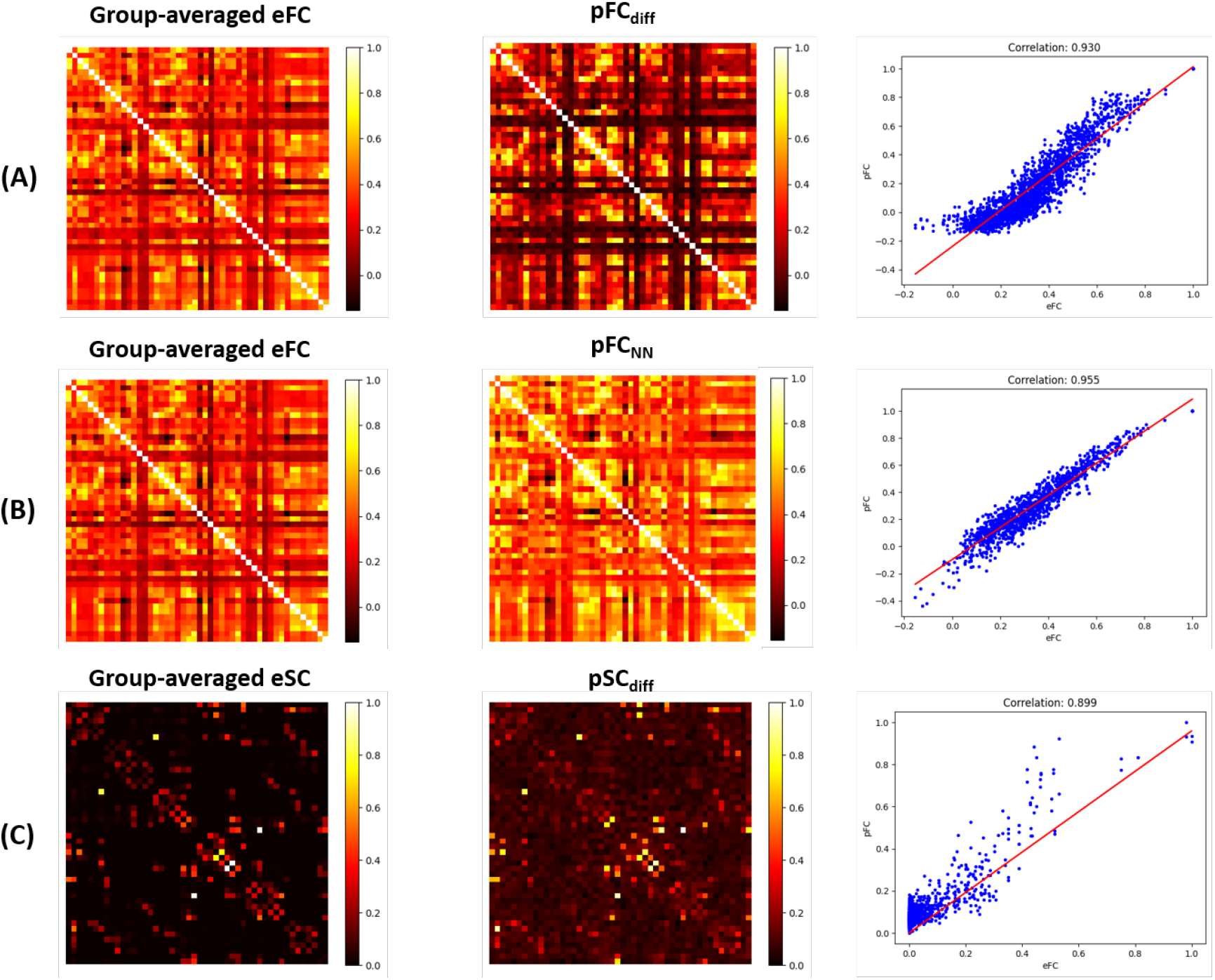
Group-averaged results for (A) UNIT-DDPM functional Connectivity. The leftmost graph is a heatmap of the group-averaged empirical functional connectivity matrix, middle graph is the group-averaged pFC predicted by the UNIT-DDPM and the right most graph is a scatter plot showing the correlation between group-averaged eFC and pFC_diff_ ; (B) Multi-layer perceptron. The figure in the left represents the heatmap of group-averaged eFC, the figure in the middle represents the group-averaged pFC predicted by the multi-layer perceptron and the figure to the right represents the scatter plot of the correlation group-averaged eFC and pFC_NN_; (C) UNIT-DDPM structural connectivity. The leftmost figure is the heatmap of group-averaged eSC, the figure in the middle is the heatmap of the group-averaged structural connectivity matrix predicted by UNIT-DDPM (pSC_diff_) and the rightmost figure is the scatter plot showing the correlation between group-averaged eSC and pSC_diff_.

Even though the FC matrices predicted by the UNIT-DDPM are more accurate on average compared to the MLP model, the difference between the intra-individual prediction accuracy of the two models is not numerically significant. Also, the overlapping of intra-individual correlation coefficients of pFC-eFCdiff within the 75th percentile of pFC-eFCNN does not rule out the null hypothesis that the difference is completely random and is of no statistical significance. To test the hypothesis that the results of UNIT-DDPM are significant better compared to the MLP model we perform a statistical test using the bootstrap method.

The basic idea of the using the bootstrap method is to make inference about the difference in average intra-individual prediction accuracy between the UNIT-DDPM and the MLP model using independent sampling with replacement from existing sample data of the same size. The results of the bootstrap method as seen in Figure 5 (C) suggests that the observed difference of 0.062 is statistically significant with a p-value of 0.0016.

## 4 Discussions

Supervised learning approaches model the SC-FC relationship by learning the unidirectional mapping between structural and functional connectomes. These models provide static snapshots of SC-FC relationship, predicting a single FC for a given SC. However, a single SC is expected to produce a range of functional networks across time, individuals and context. Therefore, supervised learning approaches cannot be considered accurate representations of this relationship. In this study, we explored an approach to infer the joint distribution of SC and FC. This provides a means to characterize the inherent variability of the SC-FC relationship. Also, we found that individual structural and functional networks are significantly more coupled than previous estimates. Interestingly, we found that significant improvements in prediction accuracy can be achieved even in the absence of SC-FC pairs.

**Figure 7.**
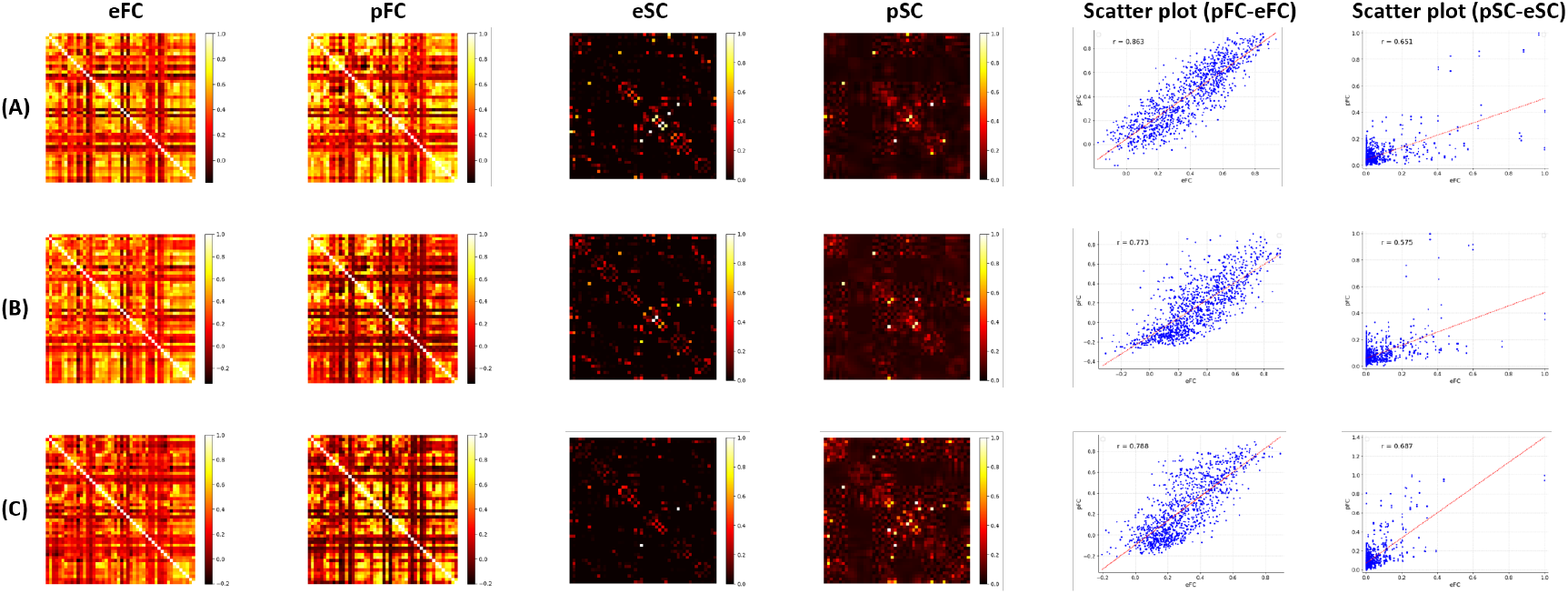
Predicting function from structure and vice versa in the human connectome using UNIT-DDPM for three representative individuals in the test set. (A) eFC, (B) pFC_diff_, (C) eSC and (D) pSC_diff_ matrices. Each row corresponds to a separate individual. Each row or column in the matrix represents a distinct sub-cortical region in the human brain. (E) and (F) Scatter plots showing the correlation between eFC *−*pFC_diff_ and eSC*−*pSC_diff_ respectively. Each point on the scatter plot represents a distinct pair of sub-cortical regions. Both pearson correlation coefficients and regression lines are indicated.

Previous studies seeking to model the SC-FC relationship have only considered predicting FC networks from structural connectomes or vice versa due to their unidirectional nature. Using a multi-layer perceptron, Sarwar and colleagues (19) achieved a correlation between empirical and predicted functional connectivity of r = 0.9 for group-averaged networks and r = 0.55 ± 0.1 for individual networks. More recently, studies have sought to predict brain function from structural connectomes and vice versa using Reinmann Networks (21), Graph Convolutional Networks (20) and GANs (22). These studies report prediction accuracies comparable to Sarwar and colleagues. But, major differences in experimental design and methodologies makes comparison between these approaches challenging. Due to our approach of sampling from an inferred joint distribution, the present study provides bidirectional estimates of SC and FC. The high accuracy of the UNIT-DDPM model in predicting group-averaged connectivity reiterates the tight coupling between structural and functional networks reported by previous studies. In terms of predicting individual networks, the UNIT-DDPM significantly outperforms existing state-of-the-art.

The high prediction accuracies reported in this study, was achieved using unpaired domain translation. This effectively bypasses the drawbacks of supervised learning, which requires empirical SC-FC pairs for training. In many cases, obtaining paired datasets can become difficult and unrealistic. The ability to accurately model the SC-FC relationship, even without paired datasets, stands as a significant advancement.

More importantly, we were able to successfully infer the joint distribution of structural and functional connectomes. This opens the possibility of identifying regions of the joint distribution with least variability in both domains, as those regions likely to provide maximum insights into the structure function relationship. Also, the ability to characterize invariant mappings in the structure-function relationship opens new possibilities for understanding the plastic response of the brain associated with various neurological disorders.

There are several limitations that require consideration. Firstly, the effectiveness of the UNIT-DDPM model with connectomes constructed using alternative pipelines and parcellation maps remains unexplored in this study. Secondly, the UNIT-DDPM model does not provide insights into the exact mechanisms governing neural activity. This lack of interpretable results is a general drawback of deep learning methods. In the future, the integration of graph topological information of brain networks with the UNIT-DDPM can be seen as possible avenues for providing interpretable results. Lastly, like many deep learning models, the complexity of UNIT-DDPM increases with the dimension of inputs, making it challenging to study connectomes with higher parcellations. To address this, alternative lightweight generative algorithms can be considered.

In summary, our study presents a novel method to characterize the inherent variability of the structure-function relationship. This approach confirms the strong correlation between brain structure and function identified in previous research. We highlight the utility of unpaired domain translation in model training, which bypasses the need for paired datasets. The accuracy in predicting individual networks demonstrates effectiveness of our approach in capturing the heterogeneity of the structure-function relationship. In the future, unpaired domain translation can be used study the changes in temporal variability associated with neurophysiological disorders.

## Acknowledgement

Data were provided (in part) by the Human Connectome Project, WU-Minn Consortium (Principal Investigators: David Van Essen and Kamil Ugurbil; 1U54MH091657) funded by the 16 NIH Institutes and Centres that support the NIH Blueprint for Neuroscience Research; and by the McDonnell Centre for Systems Neuroscience at Washington University.

